# Collection of triatomines from sylvatic habitats by a Chagas-positive, trained scent-detection canine in Texas, USA

**DOI:** 10.1101/2022.09.14.508062

**Authors:** Devin M. Christopher, Rachel Curtis-Robles, Gabriel L. Hamer, Justin Bejcek, Ashley B. Saunders, Walter D. Roachell, Thomas Leo Cropper, Sarah A. Hamer

## Abstract

**Background:** Triatomine insects, vectors of the etiologic agent of Chagas disease (*Trypanosoma cruzi*), are challenging to locate in sylvatic habitats. Collection techniques used in the United States often rely on methods to intercept seasonally dispersing adults or on community scientists’ encounters. Neither method is suited for detecting nidicolous habitats, which is important for vector control. Furthermore, manual inspection of suspected harborages is unlikely to reveal novel locations and host associations. Similar to a team that used a trained dog to detect sylvatic triatomines in Paraguay, we worked with a trained scent detection dog to detect sylvatic locations of triatomines across Texas.

**Principle methodology/Findings:** Ziza, a 3-year-old German Shorthaired Pointer previously naturally infected with *T. cruzi*, was trained to detect triatomines. Over the course of 6 weeks in the fall of 2017, the dog and her handler visited 18 sites across Texas. The dog detected 60 triatomines at 7 locations; an additional 50 triatomines were contemporaneously collected at these sites without the assistance of the dog. Approximately 0.98 triatomines per hour were found when only humans were conducting searches; when working with the dog, approximately 1.71 triatomines per hour were found. In total, 3 adults and 107 nymphs of four species (*Triatoma gerstaeckeri, Triatoma protracta, Triatoma sanguisuga*, and *Triatoma indictiva*) were collected. PCR testing of a subset revealed *T. cruzi* infection, including DTUs TcI and TcIV, in 28% of nymphs (n=100) and 66% of adults (n=3). Blood meal analysis of a subset of triatomines (n=5) revealed feeding on Virginia opossum (*Didelphis virginiana*), Southern plains woodrat (*Neotoma micropus*) and eastern cottontail (*Sylvilagus floridanus*).

**Conclusion/Significance:** A trained scent detection dog enhanced triatomine detections in sylvatic habitats. This approach is especially effective at detecting triatomine nymphs and nidicolous locations. With new knowledge of specific sylvatic habitats and key hosts fed upon by triatomines, there are opportunities for continued exploration of novel vector control methods to block the transmission of *T. cruzi* to humans and domestic animals.

**Author summary:** Triatomine insects, also known as ‘kissing bugs,’ are vectors of *Trypanosoma cruzi*, the parasite that causes Chagas disease in humans, dogs, and other mammals. Triatomines are found throughout the Americas, but little is known about where these blood-sucking insects spend their early life stages. Knowing more is important to vector control initiatives aimed at interrupting Chagas disease transmission. Scent detection dogs have been trained to detect many pests, including one study of a triatomine detection dog in South America. In this study, we used a dog to detect triatomines in their natural environments throughout Texas. Over 6 weeks, the dog identified 60 triatomines at 7 different locations; an additional 50 triatomines were collected without the dog’s assistance. Nearly twice as many triatomines were collected per hour when the dog was searching as compared to only humans searching. Of all the triatomine nymphs collected, 28% were positive for *T. cruzi*, and the blood meal analysis revealed kissing bugs had fed on Southern plains woodrat, opossum, and eastern cottontail. This study outlines a strategy that can be replicated in the United States to enhance the detection and control of habitats where triatomines spend their early life stages.

## Introduction

Chagas disease (American trypanosomiasis) is caused by the vector-borne protozoan parasite *Trypanosoma cruzi*. Approximately 5.7 million people across Latin America [1] and 300,000 people in the United States [2] are estimated to have Chagas disease. Infection also occurs in a variety of mammalian species in the United States [3], including dogs, which are burdened with disease throughout the southern United States [4–10]. The triatomine insect vectors of *T. cruzi* are found throughout the Americas, including the southern half of the US, where 11 species have been documented [11]. Limited Chagas disease treatments leave vector control as a primary means of reducing the burden of disease. Although many countries have executed successful intra-country and inter-country vector control programs throughout the past several decades [12,13], elucidating the sylvatic sources of triatomine populations remains a critical component to understanding triatomine natural history and further improving vector control efforts.

In the southern United States, triatomines exist mainly in the sylvatic environment, where they transmit *T. cruzi* among wildlife reservoirs [3]. They occasionally disperse to domestic and peridomestic environments, where they pose a risk of transmission to humans and domestic animals. For example, triatomines have recently been collected by researchers and community scientists from houses, dog kennels, porches, and garages [14–17]. However, these collections were mainly of adult triatomines and predominantly during the summer months when the adult insects were dispersing by flight from their nidicolous habitats. These findings reveal little about the habitats that support nymphal development. Although triatomines have been frequently collected from wood rat (*Neotoma* spp.) middens across the southwestern United States [18–25], relatively little is known about other blood meal sources triatomines may successfully and/or commonly utilize, particularly during their flightless nymphal stages. Manual searching for triatomines in sylvatic habitats is time- and labor-intensive, and success may be affected by the searcher’s familiarity with likely locations of triatomine habitat. For example, conspicuous harborage sites [8] and *Neotoma* woodrat middens may be more likely to be searched, although less conspicuous triatomine nidal sites likely exist.

Scent detection canines possess the ability to discern an incredible variety of odors and have been trained to identify diverse targets, including hidden explosives, fire accelerants, hazardous chemicals, illegal drugs, humans (search-and-rescue), drowning victims, invasive species, and endangered species [26]. Medical scent detection dogs are increasingly used for pre-screening of humans for infectious agents (e.g., SARS-CoV-2 [27,28]; malaria [29]; *Clostridium dificile* [30,31]) or chronic medical conditions (e.g., diabetes alert dogs [32]). Detection canines have also been trained and used for the detection of insects such as bed bugs (*Cimex lectularius*) [33,34], eastern subterranean termites (*Reticulitermes flavipes*) [35], spongy moths (*Lymantria dispar*) [36], brown marmorated stink bugs (*Halyomorpha halys*) [37], and spotted lantern fly (*Lycorma delicatula*) egg masses [38]. In addition, canines have been trained to detect primary screwworm (*Cochliomyia hominivorax*) larvae and animals infected with screwworms [39], as well as animals infected with sarcoptic mange [40], demonstrating application of detection canines for disease detection and control. Of most relevance, a study in Paraguay used a detection canine to identify sylvatic sources of *Triatoma infestans* in the Paraguayan Chaco. In this region, the sylvatic existence of *T. infestans* poses major problems to domestic control if sylvatic insects serve as a source population. At sites where previous triatomine collection efforts using light traps and Noreiau traps were largely unsuccessful, the use of the trained canine resulted in collecting 70 triatomines over a 4-month period, including a novel insect haplotype, from a variety of sylvatic sources [41].

We aimed to apply the concept introduced by the Paraguay group for canine detection of triatomines in different settings across Texas. Our objectives were to (i) demonstrate that a scent detection dog could be used to detect sylvatic habitats of triatomines in the United States; (ii) collect live insects to supplement a laboratory colony of triatomines; and (iii) determine the infection prevalence, *T. cruzi* genetic strain, and blood meal hosts of collected triatomines.

## Materials and Methods

### Ethics statement

All dog handling was completed in adherence with animal use protocols approved by Texas A&M University Institutional Animal Care and Use Committee under protocol number 2015-0289.

### Site characterization

Field work was conducted in Texas, United States, where seven species of triatomines have been documented [11]. Initial triatomine scent detection training was completed in College Station, Texas. Attempts to detect triatomines took place at 18 additional sites throughout Texas (Fig 1); sites were selected from areas with known triatomine presence, based on submissions to our community science program [42] and our own field work [16]. This study was conducted in the late fall (October 17th – December 1st, 2017), outside of the peak adult triatomine flight and collection period in Texas (generally mid-April through mid-October [16,25]), when the need for non-traditional methods of triatomine collection from nidicolous habitat is necessary for successful collection.

**Fig 1.**
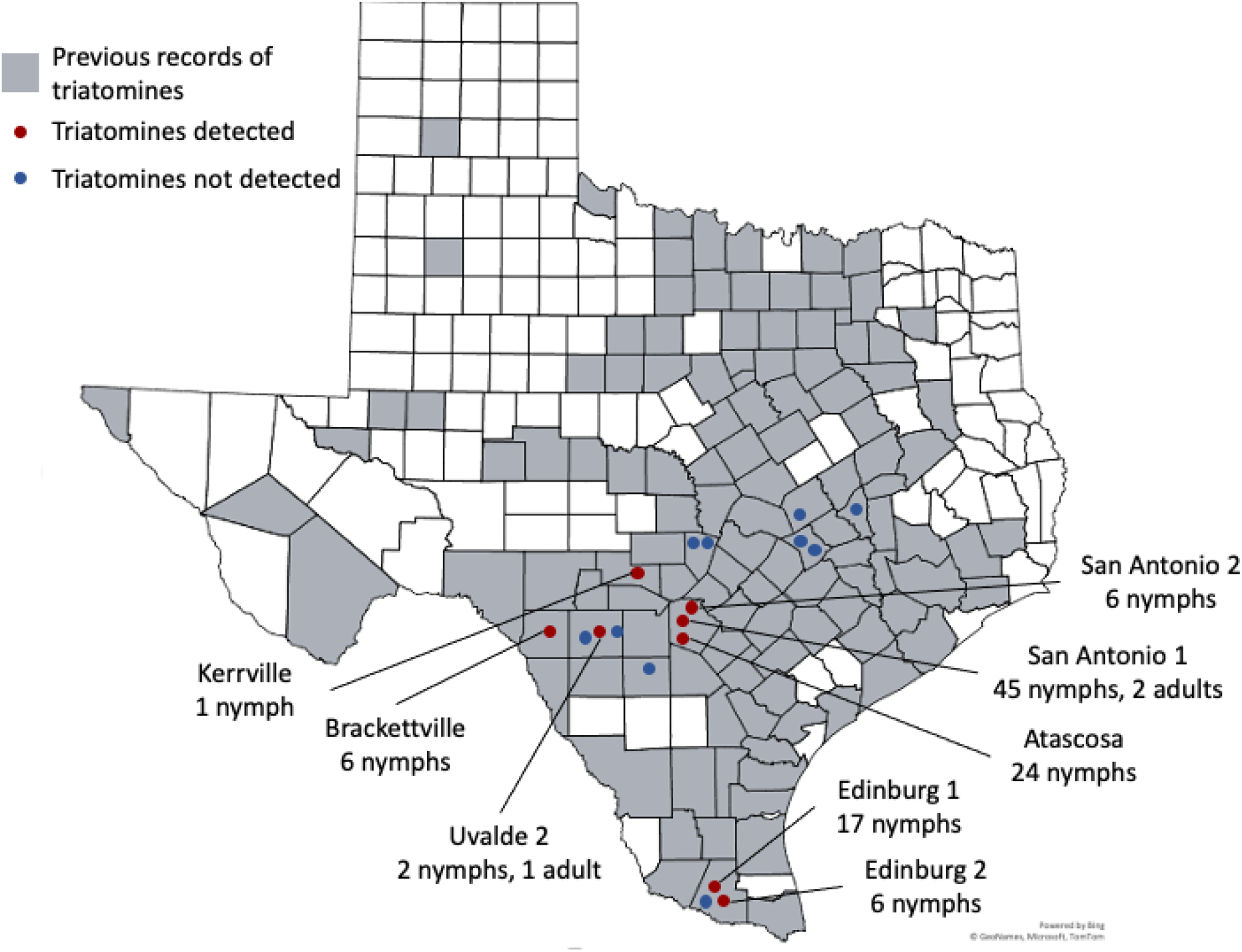
Field sites across Texas where triatomine scent detection by a trained canine were conducted. Canine scent detection efforts were conducted at locations where kissing bugs had been submitted by community scientists or collected by our field team previously. Triatomines were found at 8 of the 18 sites searched. Previous records of triatomines are based on our field collections and submission to the Kissing Bug Community Science Program [16].

### Canine health history and monitoring

As a dog trained to detect triatomines may physically contact *T. cruzi*-infected triatomines and their feces, there may be a high risk of parasite transmission to the dog. As such, we conducted our study with a previously (naturally) infected, but asymptomatic, dog.

‘Ziza,’ a German Shorthaired Pointer born in July 2014 was originally trained as an explosive detection dog by the United States government.

As part of a routine semi-annual exam in June 2016, medical records showed she tested positive via PCR for *T. cruzi* and positive for anti-*T. cruzi* antibodies on an indirect florescent antibody (IFA) test (titer value of 8192); IFA testing over the next six months remained positive. Thoracic radiographs and ECGs during these six months did not reveal any clinically significant abnormalities. Upon diagnosis with asymptomatic *T. cruzi* infection, the dog was retired from her work and was adopted several months later privately (by author DMC) in January 2017, for the purpose of training for triatomine detection. The owner noted no deviations from normal physical activity.

During this study, several tests were performed to monitor potential impacts of the existing *T. cruzi* infection on the dog’s health. Pre- and post-field work (October and December, 2017), blood was collected from the dog for testing. *T. cruzi* antibody testing was performed on serum (pre-study) or plasma (post-study) using IFA testing at the Texas Veterinary Medical Diagnostic Laboratory (College Station, TX). DNA extracted from blood buffy coat layer was subjected to real-time PCRs for *T. cruzi* DNA detection and DTU typing [43–46] using three treatments of variable DNA concentrations (neat, 2X, 1:10 dilution) to optimize chances of assigning a DTU. A 5-lead continuous read ambulatory ECG ‘Holter’ monitor (LabCorp, Burlington, NC) was applied for 48 hours. Tracings obtained from the ECG were recorded at 25mm/s using a lead configuration of V1, V2 and V5, then transferred to LabCorp for automatic analysis. Tracings were reviewed by a board-certified veterinary cardiologist (author ABS).

### Training of scent detection canine

Eight weeks before beginning her work on triatomines, the dog entered a pre-training period to review prior training and learn ‘digging’ as a secondary indication to indicate exact location upon finding. A ‘sit’ remained the dog’s official primary indication of a find. The dog was worked on a 6-meter lead.

Practice with triatomines and their scents began in October 2017. Laboratory-raised, uninfected *T. gerstaeckeri* nymphs and filter paper heavily contaminated with feces from the triatomine colonies at Texas A&M University were the primary training samples, but both live and dead *T. gerstaeckeri* adults were also used. For biocontainment, live triatomines were placed inside air-permeable, but escape-proof, containers before use in training exercises. The number of nymphs ranged from 1-10 in a single container; containers were of varied sizes and materials, such as glass, plastic, and metal, so that the dog would not associate a particular container with the triatomine scent.

The first day of training consisted primarily of imprinting activities and simple practice searches in which a contained live triatomine was presented, and the dog was rewarded with a tennis ball for sniffing the triatomine. The complete search behavior chain was taught by progressive shaping and positive reinforcement. Having previous scent detection experience, the dog quickly learned the new odor and could perform simple indoor searches before the end of the first day. The level of difficulty of the search problems was increased incrementally, until the dog could search consistently for longer than 10 minutes and reliably detect the test sample(s) in a search area at least 60 meters in width. The first day and a half, training searches were performed indoors in a closed uncluttered garage with few hiding places. During the second and third days, training searches took place in an outdoor yard with gravel, leaves, and short grass. The fourth and fifth days, training searches took place in tall grass and wooded areas. By the end of one week (a total of approximately 19 hours of the dog actively training), Ziza could reliably and quickly locate live triatomines in containers placed inside hollow logs, underneath small piles of brush or lumber, and lightly buried under dirt and leaves (to closely resemble triatomine natural habitat localities).

### Field work

After one week of training, Ziza, her handler, and at some sites, an assistant, visited 18 field sites across Texas (Fig 1). Sites were visited in October and November (2107) for two reasons: 1) weather is cooler and less stressful for the dog, and 2) dispersal of adult triatomines is minimal, so the dog would hopefully be detecting nidicolous sylvatic sources containing nymphs. Most days began with a practice problem using training samples as outlined above. The search area was then checked over by the handler and assistant prior to working the dog to identify hazards and choose the optimal search locations. Candidate search areas with obvious hazards such as broken glass, sharp metal, or dense patches of spiny plants were searched only by the human team members. The dog searched habitats including, but not limited to, woodpiles (including some piles of telephone poles, heavy (non-spiny) vegetation, downed logs, and animal nests. She was encouraged to check each of the search areas as the handler and/or assistant turned items over and created room for her to access areas of interest. When the dog displayed her primary (sit) and/or secondary (dig) indication (Fig 2A and B), the location was marked for further inspection by the handler/assistant. Searching sessions were allowed to continue so long as the dog was searching happily and did not display signs of discomfort such as heavy panting, slow movement, lying down, or when all suitable search areas were exhausted. When conditions allowed (e.g., temperature lower than 21° C, search area fully shaded, and searching primarily woody debris and logs) the dog could work as long as two hours and rest as little as 20 to 30 minutes before working again. When conditions were difficult (e.g., temperature higher than 30° C, search area in full sun, or searching primarily areas with thorny plants or other dangers) the dog could only search for 15 to 20 minutes and then given a minimum of 1 hour to rest before working again.

**Fig 2.**
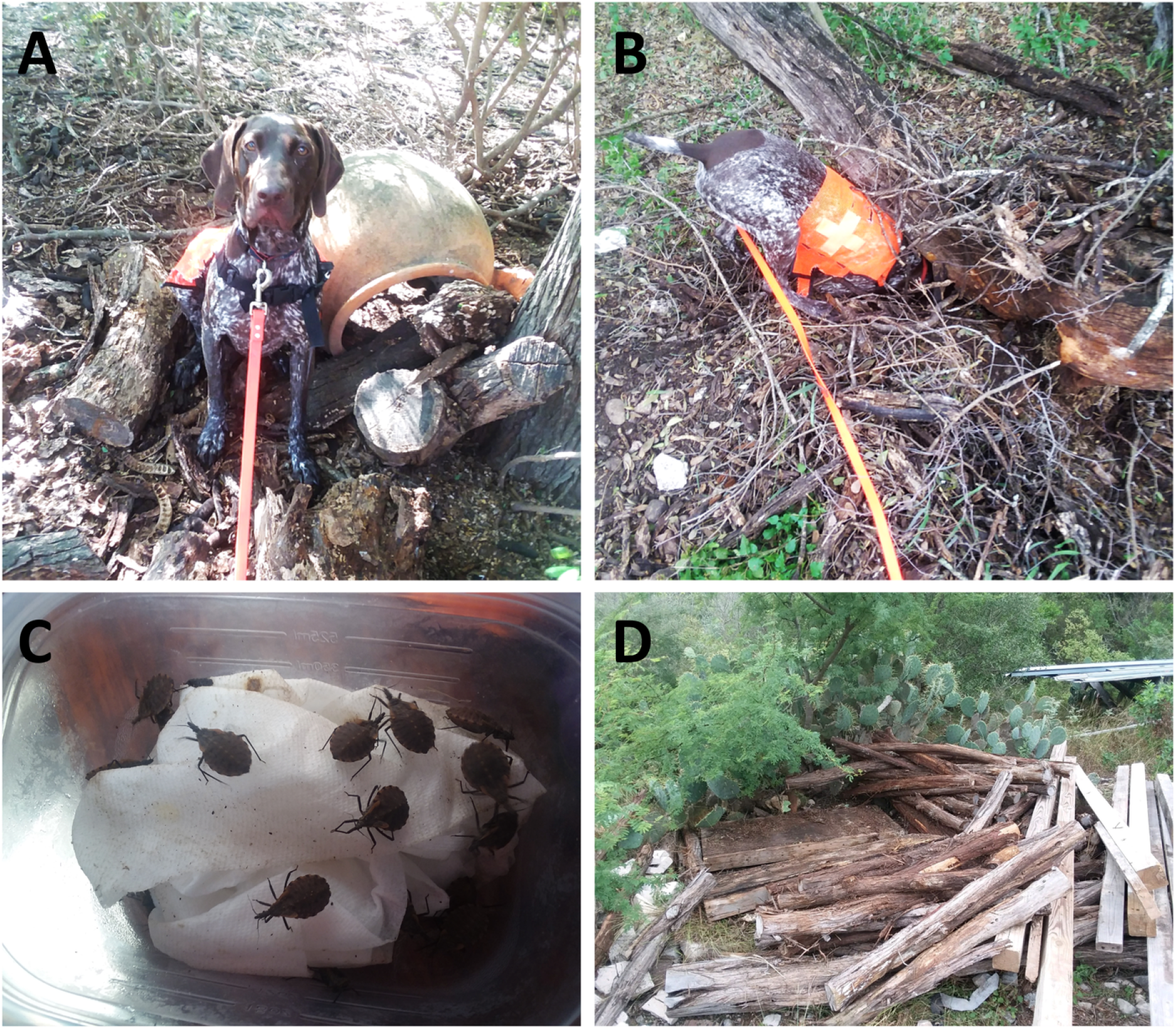
Triatomine detection canine in the field. (A) ‘Sit’ was the primary indication of a detection. (B) ‘Dig’ was the secondary indication of a detection. (C) Blood fed nymphs found at a south Texas site, in a small woodpile in the corner of a horse pen where rabbits had been seen. (D) woodpile where nymphs were found.

Triatomine findings were classified as one of three types: 1) triatomines located solely by dog, which did not require the handler or assistant to move anything in order for the dog to indicate the presence of triatomines, although the handler/assistant may have had to move the item to see and collect the triatomines; 2) triatomines located by dog with a human assist, which required the handler and/or assistant to move a large object or turn over one or more objects before the dog indicated the presence of triatomines; 3) triatomines located by human team members with no assistance by dog. Searches by humans, without the dog, were done primarily during daylight hours when it was very hot and while the dog rested, this included areas where the dog may have showed interest but had not indicated (sit and/or dig). Human searching included flipping over logs, railroad ties, and lumber/debris from piles; no intensive digging or movement of very large items was done.

### Triatomine infection with *T. cruzi* and blood meal analysis

Adult triatomines were identified to sex and species using morphologic features [47]. A subset of specimens was subjected to *T. cruzi* testing. Live specimens that entered our laboratory colony were housed individually until they generated a fecal spot on filter paper that could be tested for *T. cruzi* DNA. Specimens that did not enter the laboratory colony were prepared and dissected as previously described [16], including determination of blood meal scores (scores of 1-5 indicating no blood to large amount of blood, respectively) and blood meal analysis. DNA was extracted from fecal spot or hindgut samples (KingFisher Cell and Tissue DNA kit, Thermo Fisher Scientific, Waltham, MA). Presence of *T. cruzi* DNA was detected via a quantitative PCR using Cruzi 1/2/3 [43–45]; confirmation of infection and determination of *T. cruzi* DTU was completed using an SL-IR target probe-based quantitative PCR method [43,46]. Samples which produced a Ct value of <35 on the Cruzi 1/2/3 qPCR and were typeable on the SL-IR qPCR were considered positive for *T. cruzi* infection. Each PCR included negative controls, as well as positive controls of DNA extracted from *T. cruzi* Sylvio X10 CL4 (ATCC 50800, American Type Culture Collection [ATCC], Manassas, VA; DTU TcI), *T. cruzi*-positive (DTU TcIV) *T. sanguisuga* collected from Lee County, Texas, and *T. cruzi* Y strain (ATCC 50832, ATCC; DTU TcII).

A subset of triatomines was processed to determine sources of recent blood meals. Hindgut DNA was subjected to PCR using previously published ‘herp’ primers [48,49], direct Sanger sequenced, and compared to existing sequences in GenBank using BLAST as previously described [50]. Negative controls (water) and a positive control of DNA extracted from white-tailed deer (*Odocoileus virginianus*) heart tissue were included in each PCR batch. A blood meal identification attempt was considered successful when there was a match with ≥98% BLAST identity in the GenBank database.

## Results

### Canine health monitoring

The scent detection canine blood samples were collected immediately before (October 2017) and after (December 2017) the field work. The anti-*T. cruzi* IFA titer value prior to field work was 1280 and after field work was 320. A blood sample collected prior to fieldwork had a Ct value of 29 on the Cruzi 1/2/3 PCR, but the DTU was unable to be determined using the SL-IR PCR. After field work, the dog was PCR-negative. A 48-hour Holter monitor placed during the final week of this project documented a sinus arrhythmia with an average heart rate of 82 beats/min and one couplet of ventricular premature contractions, as well as periods of sinus arrest with the maximum lasting 5.3 seconds.

### Triatomine collections

The dog/handler team visited a total of 18 field sites across Texas over a period of six weeks during October and November 2017. Triatomines were collected at 8 sites (Fig 1). An estimated 86 hours were spent actively searching, including 35 hours of dog and handler working together, and 51 hours of humans searching without dog during the day. Search time did not include time spent getting the dog out and ready, putting the dog away, breaks, or gathering and placing bugs into the containers after finding them.

A total of 110 live triatomines were collected, including 107 nymphs (97.2%) and 3 adults (2.7%; two *T. protracta* and one *T. sanguisuga*). There were 3 instances in which a single triatomine was detected; all other detections were of triatomines in groups: 2 groups of 2 triatomines each, 2 groups of 3 triatomines, 3 groups of 6 triatomines, 1 group of 10 triatomines, 2 groups of 14 triatomines, 1 group of 17 triatomines, and 1 group of 24 triatomines. A total of 51 (46%) of the triatomines from 5 sites were located by the dog without assistance from other team members; and a total of 9 (8%) triatomines from 2 sites were located by the dog with human assistance. A total of 50 (45%) specimens from 3 sites were located by human team members only (Table 1). The majority of the human-only collections came from a single site (San Antonio site 1) from harborage locations previously known to contain triatomines. All other specimens collected by the dog and/or humans were collected from newly discovered harborage locations. We found approximately 0.98 triatomines per hour when only humans were conducting searches, even at the site (San Antonio 1) that was known to harbor triatomines. When working with the dog, we found approximately 1.71 triatomines per hour in a wide variety of microhabitats.

**Table 1.** Search success by detection type and calculation of specimens found per hours searched. Detections were classified as: 1) triatomines located solely by dog; 2) triatomines located by dog with a human assist; 3) triatomines located by human team members. Searching hours were grouped by those during which the dog was active and those during which only humans were searching (generally while the dog was resting).

| Detection type | Nymphs found | Adults found | Searching hours | Triatomines per hours searched |
| --- | --- | --- | --- | --- |
| Dog only | 50 | 1 *T. protracta* | 35 | 1.71 |
| Dog with human assist | 8 | 1 *T. protracta* |  |  |
| Humans only | 49 | 1 *T. sanguisuga* | 51 | 0.98 |

Searches were performed in habitats with a diversity of potential triatomine locations, including wood piles, stacked bricks, sheet metal on ground, stone, cactus, along walls, inside sheds, underneath potted plants, and in farming equipment. However, wood piles (specifically natural wood or railroad ties but not treated lumber) comprised the predominant location type in which nymphs were found (99%; 106 of 107 nymphs were found in these habitats). One exception was a single nymph found by a human in a trash pile consisting primarily of metal and plastic. All three adult specimens were found with nymphs (all found in woodpiles). Many specimens were found on the underside of a log or railroad tie, usually at the bottom of a moderate to large sized pile. Nymphs were typically found in shady areas with moist soil and other invertebrates, such as striped bark scorpions (*Centruroides vittatus*), sand cockroaches (*Arenivaga* sp.), ant lions (*Myrmeleon* spp.), and pillbugs (*Armadillidium* sp., *Cubaris* spp.). Specimens were often found in or near sites with dog kennels or evidence of rodents (feces and nesting materials). On more than one occasion, toads (*Bufo* spp.) were found in areas where triatomine nymphs and rodent feces were also found.

Of the 107 nymphs collected, several, but not all, were individually tracked over time after being placed alive in the TAMU kissing bug colony. Thirty-two nymphs collected from 10/31/17 and 11/1/17 at the San Antonio 1 site were combined in the same colony tub; all except one eventually molted to adults morphologically identified as *T. gerstaeckeri*, while the remaining nymph molted into an adult morphologically identified as *T. indictiva*. Fourteen nymphs collected 11/17/17 from the Edinburgh 1 site were combined in the same colony tub, and all eventually molted to adults identified as *T. gerstaeckeri*.

Of the 107 nymphs collected, 90 were tested for *T. cruzi* infection, and 13 (14.4%) were positive for *T. cruzi;* all 13 were infected with *T. cruzi* DTU TcI (Table 2). Of the three adult triatomines collected, 2 were *T. protracta* and 1 was *T. sanguisuga*. All three adult triatomines were tested, and 2 (66%) were positive for *T. cruzi* TcI, including one *T. sanguisuga* and one *T. protracta* (Table 2). A subset of 5 specimens (4 nymphs and the one *T. cruzi*-positive adult *T. protracta*) were sacrificed and subjected to dissection, blood meal scoring, and blood meal analysis, which revealed sequences with ≥98% BLAST matches to *Neotoma micropus* (Southern plains woodrat; found in 1 *T. protracta* adult, which had also tested positive for *T. cruzi* TcI, from the San Antonio 1 site), *Sylvilagus floridanus* (eastern cottontail; found in 1 nymph, which had tested negative for *T. cruzi*, from the Edinburgh 2 site), and *Didelphis virginiana* (opossum; found in 3 nymphs collected from the same Atascosa site, which had all tested positive for *T. cruzi* TcI) (Table 3).

**Table 2.** *T. cruzi* infection in triatomines collected by a scent detection canine and her human handler. From October to December 2017, a total of 110 triatomines (107 nymphs and 3 adults) were collected at 8 sites across Texas. A subset was tested for *T. cruzi* DNA from either fecal spots or hind gut dissections. Only DTU TcI was detected in the triatomines collected in this study.

| Site | Triatomines<br>found | Triatomines<br>tested | Number positive<br>(%) |
| --- | --- | --- | --- |
| Atascosa | 24 | 23 | 11 (47.8%) |
| Brackettville | 6 | 6 | 0 (0%) |
| Edinburgh 1 | 17 | 15 | 0 (0%) |
| Edinburgh 2 | 6 | 6 | 0 (0%) |
| Kerrville | 1 | 0 | NA |
| San Antonio 1 and 2<sup>a</sup> | 53 | 30 | 2 (6.7%) |
| Uvalde 2<sup>b</sup> | 3 | 3 | 2 (66.7%) |
| Total | 110 | 83 | 15 (18.1%) |

<sup>a</sup> Triatomines from these sites were combined in the colony and not followed individually. These numbers include two adults (one *T. protracta* and one *T. sanguisuga*), both of which were positive for *T. cruzi* DNA, DTU TcI.
<sup>b</sup> These numbers include one adult *T. protracta*, which was *T. cruzi* negative, and 2 nymphs that molted into *T. gerstaeckeri* adults in the colony

**Table 3.** Blood meal analysis of triatomines found by a scent detection canine. A subset of 5 triatomines were subjected to blood meal analysis to determine recent blood meal hosts. The three nymphs with opossum findings were all found in the same nest area.

| ID | Blood meal score | Species | Found by | Location | Location description | *T. cruzi* status (DTU) | Blood meal source |
| --- | --- | --- | --- | --- | --- | --- | --- |
| D+Z 017 | 3 | *T. protracta* | Dog with human assist | San Antonio 1 | Small pile of railroad ties with thorns and trees | Positive (TcI) | *Neotoma micropus* (Southern plains woodrat) |
| PS 3395A | 4 | *Triatoma* nymph | Dog | Edinburgh 2 | Pile of railroad ties in livestock pen with brush piles | Negative | *Sylvilagus floridanus* (eastern cottontail) |
| D+Z 020A | 4 | *Triatoma* nymph | Dog | Atascosa | Pile of railroad ties in a ditch | Positive (TcI) | *Didelphis virginiana* (opossum) |
| D+Z 020B | 5 | *Triatoma* nymph | Dog | Atascosa | Pile of railroad ties in a ditch | Positive (TcI) | *Didelphis virginiana* (opossum) |
| D+Z 020C | 5 | *Triatoma* nymph | Dog | Atascosa | Pile of railroad ties in a ditch | Positive (TcI) | *Didelphis virginiana* (opossum) |

## Discussion

The nidicolous, cryptic nature of nymphs makes the identification of their location a particular challenge, yet the identification of such habitats is critical for effective vector control since the adults that eventually interact with people and dogs emerge from these areas where development occurs. The use of a scent detection dog can help to achieve this objective. We trained a scent detection canine to reveal nidicolous locations of triatomines across Texas. The dog and handler team collected a total of 110 triatomines of four species across 8 locations. *T. cruzi* infection was found in 28% of nymphs and 66.6% of adult triatomines tested, congruent with vector infection prevalence estimates from our previous state-wide initiative [51]. Blood meals from Southern plains woodrat, eastern cottontail, and opossum were detected. This study expands the proof-of-concept work conducted by a Paraguay team and their triatomine detection dog, Nero, [35] to show that a trained scent detection dog can be a valuable asset for revealing nesting locations of multiple species of triatomines; in our case, nearly doubling the rate of triatomine collection compared to human-only efforts at the sites.

Due to concerns for the risk of *T. cruzi* transmission to a scent detection dog, we trained an asymptomatic Chagas positive dog. Our rapid success in training the dog was partially owed to the dog’s extensive prior training as an explosives detection dog. While Ziza repeatedly tested seropositive and PCR positive for *T. cruzi* after her initial diagnosis, she remained asymptomatic since her diagnosis in June of 2016. The dog’s positive serology and contradictory PCR results from pre- and post-field work (positive in October 2017; negative in December 2017) are reflective of the difficulty of detection of *T. cruzi* infection using PCR. Fluctuating PCR results in chronically-infected dogs have been documented as a main reason to use serology for a diagnosis [52]. Effects of reinfection on canine serology, PCR results, and disease progression have rarely been studied; one study documented progressively decreasing parasitemia and increasing antibody levels after reinfections [53]. The diagnosis, lack of symptoms, and previous detection experience made this dog an ideal candidate for triatomine detection.

The number of nymphs collected using the scent detection canine was remarkable given our research team’s experience attempting to collect triatomines from nidicolous habitat. Submissions of nymphs to community science programs in the United States are rare (two separate groups found that ~96% of submissions by community scientists were adult triatomines [16,17]), perhaps because of nymphs’ inability to fly to locations where humans are likely to encounter them and also because of their smaller and more cryptic appearance compared to the adult triatomines. Triatomines are well-documented as occurring in woodrat nests [54–56], which are distinctively built and recognizable [54,57]; using a dog helps to eliminate human visual search image and potential sampling bias and helps identify less easily-observed triatomine harborage sites including sites where other wildlife species dwell, as revealed by the blood meal analysis findings that included eastern cottontail and Virginia opossum blood.

Strikingly, and a testament to the dog’s high standard for indicating, in our training and study, the dog did not falsely indicate at any locations (i.e., every time she indicated, the human was able to find a triatomine); however, it is unknown how many times the dog may have not signaled when in fact there was a triatomine in the area. In some cases, the dog lingered at a location without indicating, and when the human returned later to search, a triatomine was found. In this study, we did not count those detections as dog or dog assisted because there was not a clear indication. We found approximately 0.98 triatomines per hour when only humans were conducting searches, driven mostly by collections as a particular site (San Antonio 1) that was known to harbor triatomines. When working with the dog, we found approximately 1.71 triatomines per hour in a wide variety of microhabitats. This is compared to our previous non-dog work and calculations of 3.8 bugs per hour when actively/destructively sampling nidicolous habitats *known* to harbor triatomines (woodrat nests in Uvalde County, Texas in July of 2014 and August of 2015) [16]. We had not previously done any other kinds of day, human-only searching in areas of unknown triatomine occupancy to which this estimate may be compared. It is important to note that the dog was able to help direct us to triatomine harborage locations that we otherwise would not have detected. This enhanced detection would be especially valuable for regions where triatomines are either not known to exist or only known to exist due to the collection of an adult specimen (e.g., [58]).

Particularly notable was the dog’s ability to detect multiple *Triatoma* species. Even though training had only used *T. gerstaeckeri* samples, the dog likely was able to detect other species, as evidenced by the findings of two adult *T. protracta* – one found by the dog only and one found by the dog with human assist. Each of the *T. protracta* samples had been found with one or two nymphs, which were likely conspecifics, but this was not confirmed. There is a paucity of information regarding how often multiple *Triatoma* species may inhabit the same nidicolous habitat, but previous studies suggest that triatomine species found in the US are typically of only one species per localized site. Therefore, there is a small but unlikely possibility that the nymphs were *T. gerstaeckeri* and detected by the dog for that scent. In addition, there was one collection that was 31 *T. gerstaeckeri* nymphs and one *T. indictiva* nymph. It is unclear from this study whether the canine would have been able to detect *T. sanguisuga*, which is a species co-occurring in areas where searches were conducted; one adult *T. sanguisuga* was found during the human only searching.

The 14% infection prevalence in nymphs we found in this work is consistent with findings of 11% from our previous work [51]. However, as more information about vector-host relationships emerges from additional collections of nymphs from nidicolous habitats, it will be interesting to see how nymphal infection prevalence is affected by hosts, including whether certain hosts are reservoir species.

We believe this is the first report of the eastern cottontail as a blood meal source. Another species of cottontail—*Sylvilagus audonbonii—* blood has been detected in triatomines, including triatomines infected with *T. cruzi* [59]. The discovery of this triatomine blood source may be due to the dog’s ability to detect nymphs in nidicolous habitats. The ability to detect blood meal sources that sustain nymphs until they can disperse as adults is a strength of this study; this information can help guide our understanding of any species that are particularly important to the early triatomine life cycle and allow for targeted interruption of the life cycle. Woodrats have previously been reported as blood meal sources in triatomines collected in Texas [55], and the findings of triatomines in woodrat nests are well-documented [57], as well as findings of *T. cruzi* in this blood meal host [60]. In our study, the triatomine with evidence of woodrat blood was infected with *T. cruzi* DTU TcI; previous work has revealed woodrats infected with DTUs TcI and TcIV [60,61]. Opossum blood has been previously detected in triatomines [62], and opossums are well-known reservoirs of *T. cruzi* [3,63], specifically DTU TcI [64–69]. As expected from this DTU-host association, and the nymphs in this study with evidence of opossum blood were infected with *T. cruzi* DTU TcI.

Of note, many of the triatomines in this study were collected from under piles of wood or railroad ties. These piles likely serve as locations where small prey animals can nest safely away from predators. The semi-protected areas under the wood also likely allowed for triatomine scent to pool and be more detectable to the dog. It may be that groups of nymphs are more easily detected than single adults moving through an area because of the scent pool that forms around the cluster of nymphs.

Multiple hazards were encountered during this field work that should be considered when planning to conduct canine scent detection work. Most relevant hazards were the sun and heat, venomous snakes, venomous arthropods, and prickly pear cactus (*Opuntia* spp., the spines of which may pose a physical hazard). Piles of scrap and sheet metal, old lumber with exposed nails, broken glass, barbed wire, and exposed metal fencing were also commonly encountered. To mitigate potential hazards, the dog was always worked on lead. A two-person team was more effective than the dog handler working alone with the dog, as handler collection of triatomines was difficult when the dog was actively excited about a detection. For personal protection, the handler wore long pants, a long-sleeved shirt, thick-soled boots, snake guards, a hat, and gloves; the dog wore a snake proof vest and a brightly colored vest, as rattlesnakes were a danger and work was conducted in areas with active hunting. Handlers should speak with their veterinarian about whether a rattlesnake vaccine is appropriate pre-exposure prophylaxis. Working during the heat of the day when temperatures exceed 27°C (80°F) in the direct sun was avoided whenever possible. A variety of potential distractions – including animals, animal feces, gunshots, and swimming pools – can be anticipated during field work and should be addressed during dog training. During fieldwork, we realized that a ‘sit at source’ indication was not the best indication for a dog doing this type of work. In many cases, the dog was unable to sit when making a detection because most detections were on the top or side of large wood piles where debris posed a sliding or otherwise unstable hazard. A bark or focused indication may be a better choice. In contrast to ‘sit at source’, digging was useful as a secondary indication.

In addition to careful consideration of local safety issues, others interested in training a canine for triatomine detection will find it most fruitful to work with an experienced scent-detection dog and skilled trainer. Ziza’s background made her an ideal candidate, as 19 hours of training were sufficient for her to recognize and signal detections of triatomines. Scent detection canines can be cost-prohibitive to include in many kinds of studies [70]. Both the method of training and the nature of the task may result in nuances in how the dog signals. Continuous training of the dog/human pair, particularly blinded training and a handler’s attentiveness to nuances in the dog’s signaling is key to the success of this method.

Although triatomines occur in 30 states in the U.S. [71], many states have relatively few documented occurrences of triatomines. Local public health agencies and triatomine researchers frequently attempt to collect triatomines from areas where human or animal cases of Chagas have occurred or where locations have been made in the past. Often considerable search effort can yield low success rates [72], and it is likely that many search efforts for triatomines that never yield positives remain unpublished. Having a scent detection canine trained specifically on triatomines would be a valuable tool in many contexts in the U.S. and beyond. Detection of triatomine nymphs in sylvatic nidicolous locations was successful using a trained scent detection dog, and future work can help to elucidate key areas for vector control for disease prevention.

## Acknowledgements

We thank Bret Nash, Italo Zecca, Edwin Valdez, Ester Carbajal, and Dr. Theresa Casey for assistance in the field. We are grateful to the property owners who allowed our team on their properties: K. and E. Avendano, Dr. J. Barnes and K. Miller Barnes, R. and B. Beall, H. Brown, M. Leonard and B. McMillian, A.V. Millard, E. Muir-Chamberlain, B. Ogea, A. Parchman, A. and B. Parker, M. Schumann, and others. Lisa Auckland provided laboratory support. Alyssa Meyers, Jillian Wormington, and Keswick Killets provided triatomine colony support. We dedicate this study to Ziza who passed away suddenly, unrelated to Chagas disease, in 2021.

## Author Contributions

Conceptualization: DMC, TLC, RC-R, GLH, SAH. Data curation: DMC, RC-R, SAH. Formal analysis: DMC, RC-R, ABS. Funding acquisition: RC-R, GLH, SAH, ABS. Investigation: DMC, TLC, RC-R, SAH, WDR. Methodology: JB, DMC, RC-R, GLH, SAH, WDR, ABS. Project administration: DMC, RC-R, SAH. Resources: JB, DMC, RC-R, GLH, SAH, ABS. Software: DMC, RC-R, SAH, ABS. Supervision: SAH. Validation: DMC, RC-R, GLH, SAH, ABS. Visualization: DMC, RC-R, SAH. Writing – original draft: DMC, RC-R. Writing – review & editing: JB, DMC, TLC, RC-R, GLH, SAH, WDR, ABS.

